# Design and discovery of ‘tug-of-war’ riboswitches

**DOI:** 10.64898/2026.06.17.733018

**Authors:** David Z. Bushhouse, Jiayu Fu, Julius B. Lucks

## Abstract

Riboswitches are compact regulatory RNAs that modulate gene expression by switching between mutually exclusive structures in response to binding diverse signalling molecules. Riboswitches that regulate transcription have been shown to function through sequence overlap between their ligand binding aptamer domain and their regulatory expression platform, a constraint that limits their engineering for biotechnological applications. Here we describe a new, modular, transcriptional riboswitch architecture, termed “tug-of-war” (TOW), that removes this constraint, based on the discovery that ligand-stabilized RNA structures positioned immediately upstream of intrinsic terminators suppress termination. Using diverse aptamers, we create four distinct TOW riboswitches by pairing aptamer domains with terminators, and discover design principles for tuning their function. We demonstrate that upstream secondary and tertiary RNA structures generally inhibit intrinsic termination, consistent with a model of TOW antitermination dependant upon steric competition for occupancy of the RNA polymerase exit channel. Finally, we discover the TOW architecture to be present in nature through bioinformatic analysis and experimental validation, finding that ∼10% of natural ZTP riboswitches – nearly all in gram-positive bacteria – employ the TOW architecture. Together, these findings define a new mechanism of nascent RNA-mediated antitermination that may be useful for searching for riboswitches in other domains of life, and establish TOW riboswitches as a highly modular platform for engineering transcriptional RNA regulators.

## INTRODUCTION

Riboswitches are cis-regulatory non-coding mRNA elements that modulate expression of associated coding sequences in response to binding of a cognate ligand (1). The ligands sensed by riboswitches (e.g. ions, metabolites, amino acids, nucleotide derivatives, tRNAs) and the regulatory mechanisms they employ (e.g. transcription termination, translation initiation, RNA degradation control, splicing modulation) are highly diverse (2, 3). Riboswitches regulate through two domains: a highly conserved aptamer domain (AD) that binds the ligand, and a downstream gene-regulatory expression platform (EP) that converts ligand binding into a gene expression outcome (4). Because many riboswitches regulate essential metabolic genes in pathogenic bacteria, riboswitches have attracted interest as targets for antimicrobial therapeutics (5). In addition, both natural and synthetic riboswitches have been widely used within biosensors, conditional expression systems, and synthetic gene circuits (6–9). As compact and dynamic RNA molecules, riboswitches have also served as powerful model systems for understanding RNA folding and RNA-small molecule interactions (10–14).

All known riboswitches work through the ability of the RNA sequence to adopt mutually exclusive folds: in the absence of ligand the EP can fold into a form that blocks or allows gene expression, while ligand binding stabilizes the AD, preventing EP folding and promoting the opposite outcome (15). This mechanism requires sequence overlap between the AD and EP that forces a structural competition for the shared sequence region. For example, in riboswitches that regulate transcription, often the AD contains a portion of the RNA transcriptional terminator hairpin that is required to form within the RNA polymerase (RNAP) exit channel to destabilize the paused transcription complex at the termination polyU site (16–19). Riboswitches that regulate transcription can also feature an EP with a central antiterminator structure that overlaps with both the AD and the terminator hairpin, inverting the regulatory logic (20–25). In either case, direct structural competition between alternative RNA folds underlies the switching mechanism. In many transcriptional riboswitches the structural competition between AD and EP happens dynamically during cotranscriptional folding (11, 25, 26). The necessity of an AD/EP sequence overlap driven by dynamic RNA folding has severely complicated engineering of transcriptional riboswitches as changes to the EP need to be compensated by changes in the AD which are often highly conserved (6, 27, 28).

While many transcriptional riboswitches feature competitive folding pathways with overlapping AD and EP domains, it is not known if this is the only way riboswitches can control intrinsic termination. For example, antitermination factors can act by allostericly modulating RNAP to mitigate the conformational changes normally induced at the polyU pause (e.g. putL RNA (29, 30)), or by narrowing of the RNAP RNA exit channel to sterically occlude terminator hairpin formation (e.g. Qλ with NusA (31, 32)). This raises the possibility of other RNA-based mechanisms to occlude terminator formation. Recently, groundbreaking studies elucidated the structural role of RNA hairpin formation in facilitating RNAP pausing and intrinsic termination (33, 34). While these studies did not include the upstream 5’ RNA context, modeling revealed a putative ssRNA exit groove in the side of the RNA exit channel (34). We hypothesized that the steric constraint of this ssRNA groove would force the RNA terminator hairpin to compete for exit channel occupancy with RNA structures upstream of the terminator. In such a model, the sterically constrained context of the exit channel would force a ‘tug-of-war’ (TOW) between the terminator hairpin and the upstream structural element, mediating sequence-independent structural competition.

Here we validate this hypothesis, and create an entirely new class of modular, synthetic TOW riboswitches that do not rely on AD/EP sequence overlap to function. We began by placing ligand-stabilized riboswitch ADs immediately upstream of various terminators, demonstrating functional switch designs. We also showed that the functional properties of these riboswitches can be easily programmed by exchanging terminator hairpins, highlighting the generality of this discovery. We also discovered additional design principles, including the need to balance the relative folding stability of the upstream structure with regard to the terminator hairpin, and the length of the spacer separating the two elements. Transcription assay results generally supported our hypothesis and revealed a TOW termination mechanism solely dependent on RNA structures upstream of the terminator, although the function of some designs appear potentially reliant on additional cellular transcriptional machinery. Finally, we performed bioinformatic sequence analysis of natural riboswitch EPs, revealing that the TOW architecture is used by natural ZTP riboswitches in Gram-positive bacteria. Validation of the function of 22 natural TOW candidates in cellular gene expression assays in *B. subtilis* and *E. coli* demonstrates that this mechanism is general across at least two bacterial phyla. Taken together, our results unveil a new mechanism of nascent RNA structure-induced transcription antitermination, that coupled with ligand binding aptamers, creates a newly discovered riboswitch mechanism. We anticipate the principles of TOW riboswitches to uncover new aspects of nascent RNA folding and function in other processes that are closely coupled to transcription, and to create a new platform for engineering RNA-based sensors and feedback mechanisms for a wide range of biotechnology applications.

## RESULTS

### ‘Tug-of-war’ riboswitches function with uncoupled aptamers and expression platforms

We first sought to test our hypothesis that placing a riboswitch AD immediately upstream of an intrinsic terminator would enable inducible transcription antitermination. To do so, we designed and functionally characterized a panel of 12 synthetic ‘tug-of-war’ (TOW) riboswitches using a modified AD from the *Clostridium beijerinckii pfl (Cbe pfl)* riboswitch that binds to ZTP, a bacterial alarmone that signals deficiencies in purine biosynthesis (18). In this system, the ligand-stabilized AD forms a robust tertiary structure in the immediate vicinity of the 3’ terminus (Fig. 1A), which we hypothesized could sterically occlude folding of the downstream terminator sequence within the RNAP exit channel. Riboswitches were constructed by placing intrinsic terminators of varying stabilities, quantified by the predicted ΔG of folding, immediately downstream of the AD, separated only by a single nucleotide spacer (Fig. 1B, S1A). The synthetic riboswitch panel was tested for ligand-inducible regulation of sfGFP expression in vivo as described in Methods. Of the 12 riboswitch designs, those with stronger hairpin stems (ΔG of < -16 kcal/mol) all exhibited >3-fold gene expression increases in response to treatment with 1 mM Z, while those with weaker terminator stems showed less response to ligand induction (Fig. 1C). Riboswitches with weaker terminators were characterized by an ‘always ON’ pattern of high gene expression in the absence of Z, suggesting that these terminators are insufficient to attenuate transcription even in the absence of Z (Fig. S1B,C). Across all terminators, both leaky expression and ligand-induced expression correlated with the predicted stability of the terminator used, with weaker terminator hairpins leading to higher levels of observed gene expression, consistent with previous reports (Fig. S1C) (35). We confirmed the transcriptional basis of these findings with IVT assays on a subset of the synthetic riboswitches (Fig. S1D).

**Figure 1:**
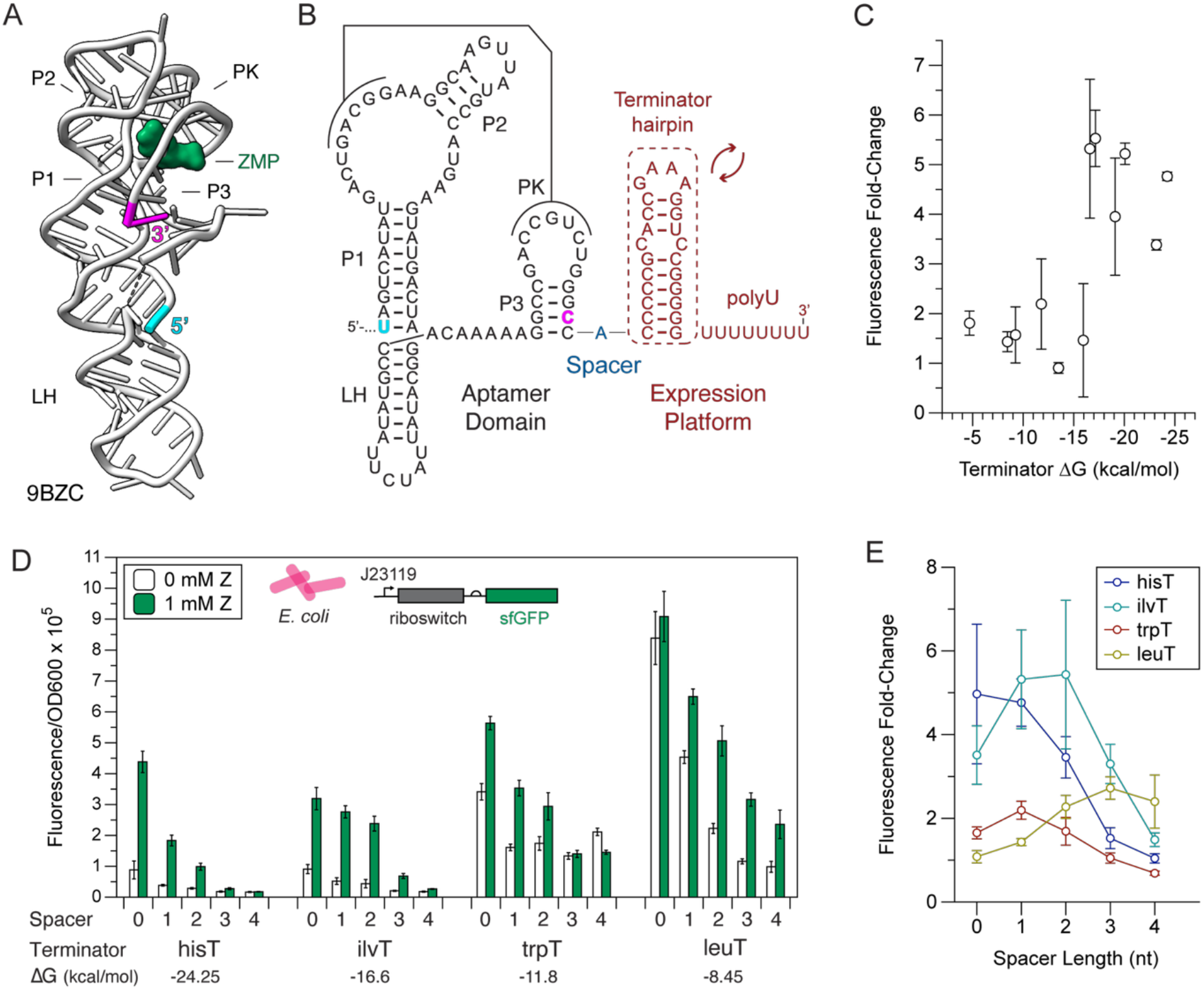
Designing and characterizing synthetic ‘tug-of-war’ ZTP riboswitches. (A) Crystal structure of ZMP-bound *Cbe pfl* riboswitch AD (PDB: 9BZC), with key structural features labelled. Cyan and magenta residues indicate 5’ and 3’ termini, respectively. (B) Secondary structure schematic of the chimeric TOW ZTP riboswitches tested in this figure. The *Cbe pfl* AD with strengthened P3 stem was placed immediately before a single nucleotide spacer, followed by an interchangeable terminator hairpin and 8 nt polyU tract. Cyan and magenta residues correspond to the structural model in (A). A panel of 12 constructs that varied in the terminator hairpin sequence were constructed (Figure S1). (C) Fluorescence fold change in a cellular gene expression assay between 0 mM Z and 1 mM Z treatment for the synthetic riboswitch panel, measured and calculated as described in Methods. The x-axis represents the predicted free energy of folding for each terminator structure (Figure S1). Error bars indicate error-propagated standard deviation of n=9 biological replicates for each construct. (D) In vivo reporter expression assays performed in *E. coli* characterizing switching function of chimeric ZTP riboswitch constructs with various terminator hairpins and polyA spacer lengths. Error bars indicate standard deviation of n=9 biological replicates. (E) Fluorescence fold change between 0 mM Z and 1 mM Z treatment for riboswitch variants in (D), measured and calculated as described in Methods. Error bars indicate error-propagated standard deviation of n=9 biological replicates.

To interrogate the mechanistic basis for ligand-induced antitermination in these synthetic riboswitches, we selected a subset of 4 riboswitches and varied the length of the spacer region from 0 to 4 nt. We hypothesized that if ligand-induced antitermination requires direct interaction between the holo-AD and RNAP, then longer spacers would provide additional slack for the terminator to fold regardless of AD ligand-binging status, leading to reduced gene expression and potentially switching. Consistent with this hypothesis, the extension of the spacer region biased all the riboswitches in the subpanel towards termination, resulting in lower readthrough gene expression in both the presence and absence of ligand (Fig. 1D, S1E). For each terminator, we found an ideal range of spacer lengths to maximize the fold change of the resulting synthetic riboswitch (Fig. 1E). For more stable terminators (ΔG of < -16 kcal/mol), shorter spacers resulted in the best performance, while longer spacers result in an ‘always OFF’ phenotype. On the other hand, the least stable terminator in the subpanel, leuT, requires longer spacers to achieve efficient switching, while shorter spacers result in an ‘always ON’ phenotype. IVT assays on a subset of this panel confirmed the general trends, though we were only able to observe functional switching for the designs with the highest fold-change in vivo, suggesting that the more complex cellular transcriptional machinery may enhance or even be required for some TOW riboswitch designs (Fig. S1E).

These results validate our initial hypothesis that aptamer domains immediately upstream of intrinsic terminators result in ligand-dependent antitermination. This finding has two immediate consequences: the potential facile construction of synthetic transcriptional riboswitches through modularly changing aptamer domains, and their straightforward functional optimization through changing the terminator stem free energy and spacer length, both of which directly control termination efficiency.

### Synthetic TOW riboswitches function with diverse aptamer domains

We next sought to extend these findings by designing additional TOW riboswitches using diverse ADs. Three ADs were selected: the natural *Bacillus cereus crcB* (*Bce crcB*) fluoride AD (Fig. 2A-C) (17), a 2-aminopurine (2-AP) sensing variant of the *Bacillus subtilis pbuE* (*Bsu pbuE*) adenine AD (36) (Fig. 2D-F), and a synthetic tetracycline AD originally developed to control translation initiation in yeast (37) (Fig. 2G-I). Each AD was paired with a panel of five terminators using three different polyA spacer lengths, for a total of 15 riboswitch designs per AD. These designs resulted in functional riboswitches (>2-fold induction) for all the selected ADs (Fig. 2, S2), and differences between the designs further demonstrated the relationship between spacer length and terminator strength in tuning riboswitch fold-change, consistent with what was observed for the ZTP AD. For example, across the different AD classes, we observe lower termination when stronger terminators or longer spacers are used, and higher expression when weaker terminators or shorter spacers are used (Fig. 2C,F,I, S2).

**Figure 2:**
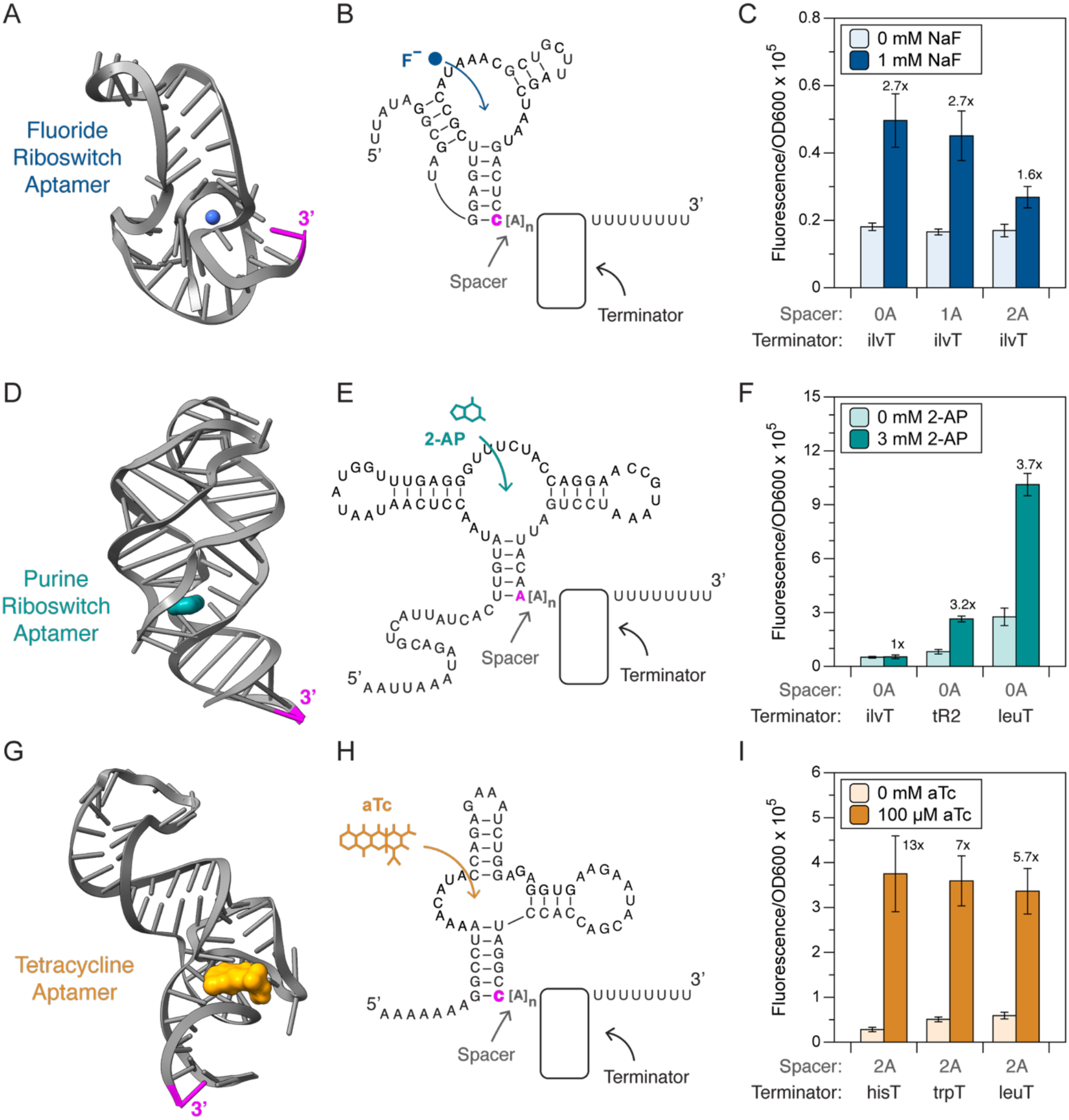
Construction of synthetic ‘plug-and-play’ riboswitches using TOW design rules. (A) Structure of the *Thermotoga petrophila crcB* fluoride riboswitch aptamer (PDB: 4ENC) (ref. (69)). Fluoride ion is indicated in blue, and the 3’ nucleotide indicated in magenta is the residue grafted onto the spacer-terminator module for each synthetic riboswitch variant. (B) Secondary structure schematic of the TOW synthetic fluoride riboswitch design, including the aptamer domain shown in (A) and a variable spacer-terminator module. (C) In vivo reporter expression characterization of synthetic TOW fluoride riboswitch designs in the presence (1 mM NaF) and absence (0 mM NaF) of cognate ligand. (D) Structure of the *Vibrio vulnificus pbuE* purine riboswitch aptamer (PDB: 5SWE) (ref. (70)). Adenine ligand is indicated in teal, and the 3’ nucleotide indicated in magenta is the residue grafted onto the spacer-terminator module for each synthetic riboswitch variant. (E) Secondary structure schematic of the TOW synthetic purine riboswitch design, including the aptamer domain shown in (D) and a variable spacer-terminator module. (F) In vivo reporter expression characterization of synthetic TOW purine riboswitch designs in the presence (3 mM 2-AP) and absence (0 mM 2-AP) of cognate ligand. (G) Structure of the tetracycline aptamer (PDB: 3EGZ) (ref. (71)). Ligand is indicated in orange, and the 3’ nucleotide indicated in magenta is the residue grafted onto the spacer-terminator module for each synthetic riboswitch variant. (H) Secondary structure schematic of the TOW synthetic tetracycline riboswitch design, including the aptamer domain shown in (G) and a variable spacer-terminator module. (I) In vivo reporter expression characterization of synthetic TOW tetracycline riboswitch designs in the presence (100 uM aTc) and absence (0 uM aTc) of cognate ligand. For C, F, I error bars indicate standard deviation of n=9 biological replicates. Fluorescence fold change for each riboswitch variant is indicated above each set of bars. Data for additional designs is available in Fig. S2.

These results also shed light on the contribution of the AD in synthetic riboswitch performance, since the best performing riboswitch for each AD required a different combination of terminator hairpin and spacer length (Fig. 2C,F,I, S2). The fluoride and purine ADs, which are apparently more labile than the tetracycline AD, performed best when paired with short spacers or weak terminators, which enabled increases in ON-state gene expression (Fig. 2C,F, S2A,B). For these ADs, long spacers and strong terminators resulted in a ‘always OFF’ phenotype, consistent with the AD providing an insufficient barrier to terminator formation. On the other hand, the tetracycline AD, which is apparently less labile than the fluoride or purine ADs, performed best with longer spacers and stronger terminators (Fig. 2I, S2C). Shorter spacers and weaker terminators resulted in elevated leaky expression for this AD, resulting in poorer fold-change.

Taken together, these findings demonstrate the generality of the TOW riboswitch mechanism, including the potential to convert any ligand-stabilized RNA domain into a functional transcriptional riboswitch. The apparent need to ‘balance’ terminator stability and spacer length for each AD also provides a rational basis for tuning TOW riboswitches using spacer length and terminator strength.

### Upstream RNA structures generally suppress transcription termination

To further interrogate the mechanistic basis for TOW antitermination, we set out to characterize the role of simpler secondary and tertiary upstream RNA structures on termination efficiency (Fig. 3A). We first tested the effect of placing the T2 gene32 pseudoknot (PT2G32), a model stable tertiary structure (38, 39), upstream of our five-terminator panel. For all terminators, we observed that the placement of the PT2G32 pseudoknot immediately upstream of the terminator reduced terminator efficiency, while extending the spacer increased termination efficiency (Fig. 3B). To determine if even simple upstream secondary structures can suppress intrinsic termination, we prepared constructs in which the strong hisT hairpin from our panel was placed upstream of the other terminators, separated by a single A spacer. To determine the role of the upstream hisT hairpin on downstream terminator efficiency, we generated a range of upstream secondary structure contexts by disrupting the bottom base pairs of hisT with C or G→A mutations (Fig. 3A). In this way, the secondary structure in the region immediately upstream of the test terminator could be linearized without altering local sequence context (Fig. 3A, S3A-D). In both cellular expression assays and in IVT reactions, the greatest effect on termination efficiency was observed with the least stable terminator, leuT, while little to no effect was observed for the more stable terminators (Fig. 3C).

**Figure 3:**
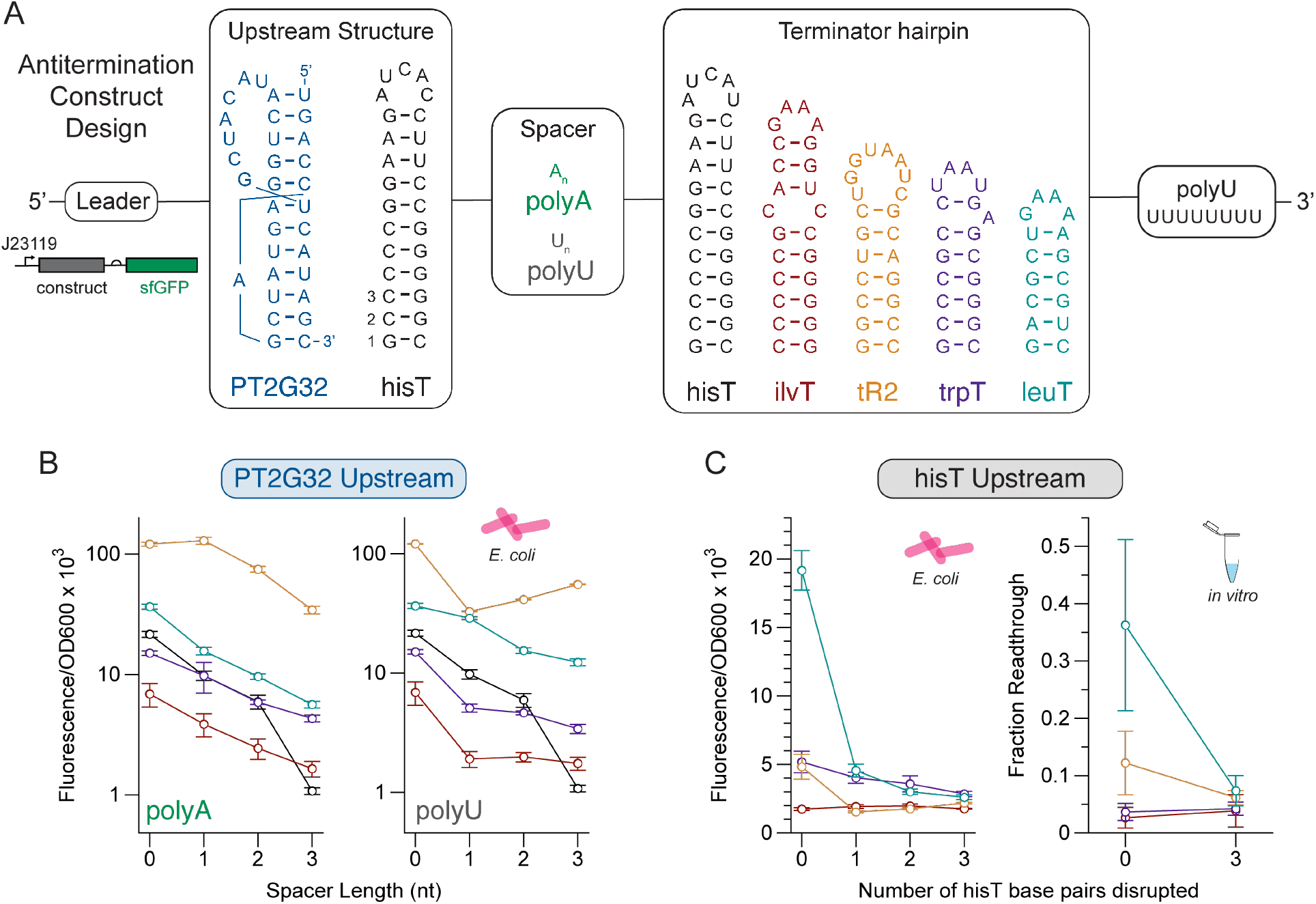
Upstream secondary and tertiary structures generally inhibit intrinsic termination. (A) Schematic of TOW antitermination constructs designs. A fixed 5’ leader sequence was placed in front of the upstream structure, either the PT2G32 pseudoknot or the hisT hairpin, followed by a variable-length polyA or polyU spacer region, one of five test terminator hairpins, and finally a 8 nt polyU tract, all upstream of an sfGFP coding sequence reporter. (B) Terminator readthrough assays performed in vivo on PY2G32 constructs illustrated in (A) with increasing polyA (left) or polyU (right) spacer length. Color coding corresponds to the terminator indicated in (A). Error bars indicate standard deviation of n=9 biological replicates. (C) Terminator readthrough assays performed in vivo (left) or in vitro (right) on hisT constructs illustrated in (A) with increasing number of C or G→A mutations in the upstream hisT hairpin at positions 1-3 labeled in (A). Color coding corresponds to the terminator indicated in (A). Error bars indicate standard deviation of n=9 (left) or n=4 (right) biological replicates.

Taken together, these findings demonstrate that relatively small upstream RNA secondary and tertiary structural elements suppress transcription termination in bacteria. While these effects are less pronounced than the antitermination effects of larger ligand-bound riboswitch ADs, we observe the same dependance on short spacer length for antitermination, pointing to a shared underlying ‘tug-of-war’ between the terminator hairpin and upstream structure. In addition, the sequence-independence of this mechanism suggests a potentially simple energetic competition between the nucleated terminator hairpin, stabilized by interactions with the RNAP exit channel, and the upstream RNA structure, mediated by the spacer region.

### Discovery of natural TOW ZTP riboswitches

Because of the relative ease with which we created synthetic riboswitches using the TOW architecture, we next asked whether natural riboswitches have evolved to employ this architecture as well. We performed a bioinformatic analysis of ZTP riboswitch sequences identified by Kim et al. (ref. (18)) and those catalogued in the Rfam database, and identified 434 unique riboswitch sequences that feature TOW architecture, i.e. with no detected overlap between aptamer and terminator sequences (Fig. 4A). Candidate sequences were identified across bacteria, but most sequences were concentrated in the *Bacilli* taxonomic class, where nearly all ZTP riboswitches possess TOW expression platforms (Fig. 4A). Interestingly, some genomes in *Paenibacillus* feature as many as four unique TOW-architecture ZTP riboswitches. Natural TOW ZTP riboswitches almost all separate the AD from the terminator with a single stranded spacer <5 nt in length (Fig. 4B), consistent with the range we observed as useful for synthetic TOW riboswitches (Fig. 1D,E). Most natural TOW riboswitches also feature terminators with ΔG < -15 kcal/mol (Fig. 4C), consistent with the functional threshold we observed for the *Cbe pfl* ZTP AD (Fig. 1C).

**Figure 4:**
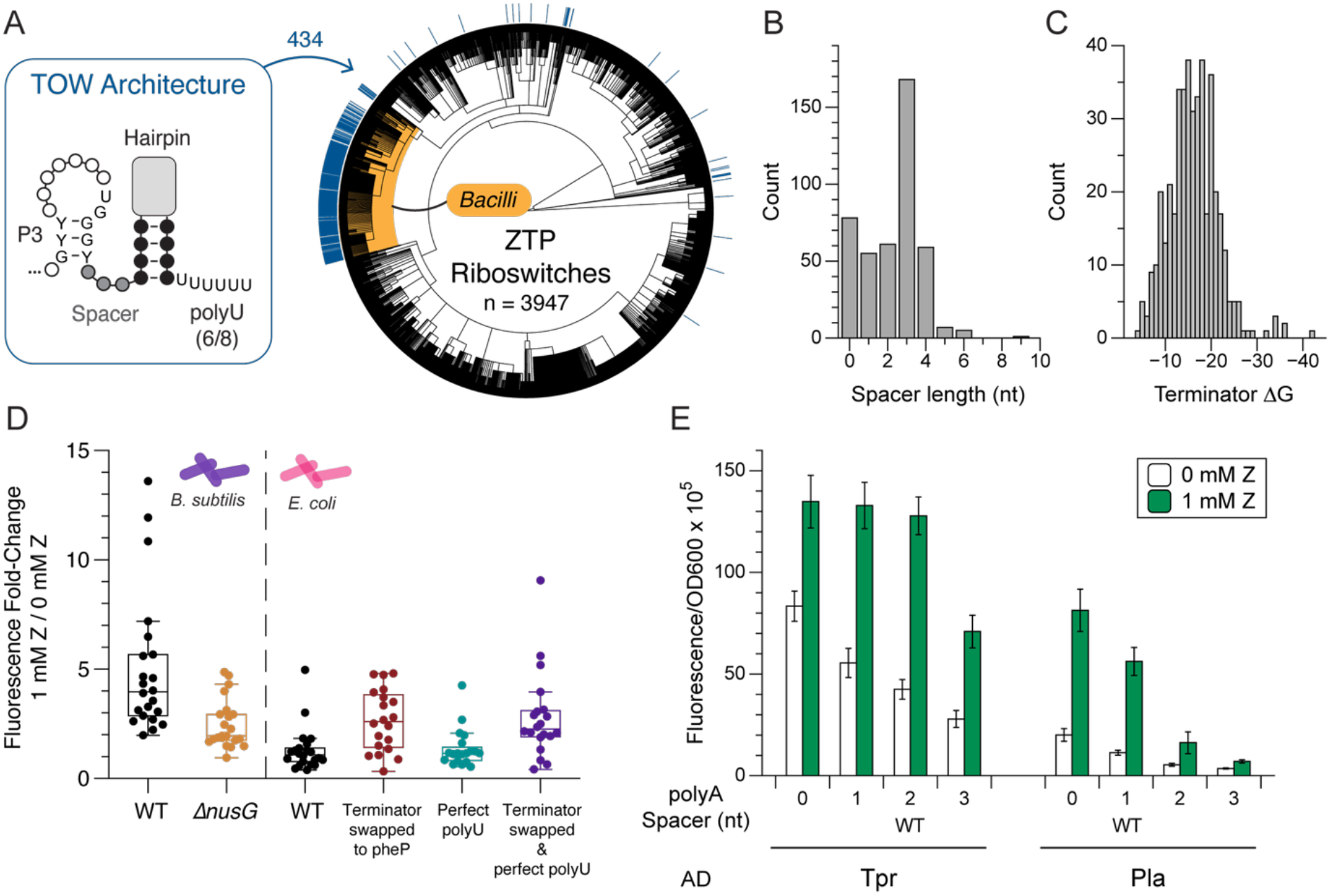
Discovery of natural ZTP riboswitches with functional TOW architecture. (A) Schematic of the motif searched for among the available identified candidate ZTP riboswitches. The TOW architecture searched for included a consensus ZTP AD P3 stem, a >= 0 nt single stranded region, a terminator hairpin, and a U-rich tract. A taxonomic tree of cataloged ZTP riboswitches, with bioinformatically identified TOW architecture labeled in blue. Most of the TOW ZTP riboswitches are found in the *Bacilli* class, shaded in yellow. (B) Histogram of spacer lengths from TOW riboswitches identified in (A). (C) Histogram of the ViennaRNA calculated free energy (ΔG) for terminator hairpins from TOW riboswitches identified in (A). (D) Box plots of the calculated fluorescence fold change between 0 mM Z and 1 mM Z treatment for riboswitches from the experimental panel. Points indicate fold change values. Data left of the dotted line was collected from experiments performed in *B. subtilis* stain PLBS727 and PLBS728 as described in Methods. Data right of the dotted line was collected from experiments performed in *E. coli* BW25113 Keio parent strain. X-axis labels on the right side of the plot indicate WT or modified sequences. (E) In vivo reporter expression assays performed in *E. coli* characterizing switching function of *Treponema primitia* ZAS-1 (Tpr) and *Paenibacillaceae bacterium* GAS479 (Pla) TOW ZTP riboswitches with various polyA spacer lengths. WT label indicates native expression platform sequence. Error bars indicate standard deviation of n=9 biological replicates.

To test discovered candidate sequences, we selected a diverse panel of 22 natural ZTP TOW riboswitch sequences, and performed in vivo characterization in both *B. subtilis* and *E. coli*. The gram-positive *Bacillus subtilis* RNAP (*Bs*RNAP) is known to respond differently to pausing and termination sites than *E. coli* RNAP (*Ec*RNAP) (40, 41), and most of the sequences in our panel are from the *Bacillus* and *Paenibacillus* genera. While nearly all of the riboswitches in our panel responded to the addition of ligand with >=2-fold increase in gene expression in *B. subtilis* (Fig. 4D,S4A), only 2 of these sequences responded to ligand induction in *E. coli* (Fig. 4D,S4B), while most of the rest displayed an “always ON” phenotype (Fig. S4B,C). The 2 functional sequences, *Tpr* and *Pla*, from *Treponema primitia* ZAS-1 and *Paenibacillaceae bacterium* GAS479, respectively, both function in vitro with *Ec*RNAP (Fig. S4D), and demonstrated the expected behavior for spacer extension, with the native spacer sequence resulting in the highest fold change for both riboswitches (Fig. 4E).

Given the discrepancy between TOW function between E. coli and B. subtilis RNAPs, we next sought to interrogate the apparent incompatibility between most of the native riboswitches in our panel and *Ec*RNAP. Specifically we attempted to restore switching function in *E. coli* by either replacing the native terminator with the strong *E. coli* terminator pheP, correcting the native U-rich tract to perfect polyU, or both (Fig. 4D). Many of the riboswitches could be made to function with >2-fold increase in gene expression by replacing the terminator with pheP, which resulted in decreased leaky expression in the absence of Z (Fig. S4C). Replacing the polyU region with perfect polyU drastically decreased gene expression in both the absence and presence of ligand (Fig. S4C), leading to an “always OFF” phenotype with poor dynamic range (Fig. 4D). The largest improvement in average fold change was achieved by replacing both the terminator and the polyU tract (Fig. 4D), which resulted in even lower leaky expression, and consistent antitermination in the presence of ligand (Fig. S4C). A selection of improved chimeric riboswitches was verified to function in vitro with *Ec*RNAP (Fig. S4D). Together, these results confirm that while most of the native intrinsic termination are insufficient to halt *Ec*RNAP, AD folding and ligand binding is maintained, and switching can be restored by swapping out EP components.

Since *E. coli* and *B. subtilis* intrinsic terminators have similar average ΔG, stem length, and polyU length (41), we wondered how the native terminators from these riboswitches are sufficient to attenuate transcription by *Bs*RNAP. Recent reports indicate that most intrinsic terminators in *B. subtilis* function with better efficiency in the presence of the transcription factor NusG (42–44). To investigate a possible role for NusG in the TOW mechanism in *B. subtilis*, we recharacterized our ZTP riboswitch panel in ΔNusG *B. subtilis*, and observed significant changes in the function of the riboswitches (Fig. 4D, S4C). Riboswitch fold change was significantly worse in the ΔNusG strain, which coincided with an increase in average leaky baseline expression but no difference in induced expression (Fig. S4C). These results indicate that NusG is likely impartant for TOW ZTP riboswitch function in *B. subtilis*, confirming earlier reports (44).

Together, these findings establish the widespread use of TOW architecture by natural ZTP riboswitches to induce antitermination in bacterial RNAP. This finding demonstrates that TOW antitermination is likely a universal feature of bacterial RNAPs that utilize intrinsic termination. Interestingly, we observed drastic differences in TOW riboswitch function between *B. subtilis* and *E. coli*, which could be ameliorated by swapping out terminator components, potentially pointing to nuanced differences between the processing of termination signals by *Bs*RNAP and *Ec*RNAP.

## DISCUSSION

In this work, we discovered an easily programmable mechanism by which nascent RNA structures block intrinsic transcription termination in bacteria, and used it to create a new class of ‘tug-of-war’ transcriptional riboswitches that we also found to occur in nature.

Our studies are based on an exploration of the mechanisms governing intrinsic transcription termination, focusing on how upstream RNA structural elements can influence terminator formation within the RNAP exit channel. Early studies of intrinsic termination demonstrated that all components of the termination site, including upstream sequence, can affect termination efficiency (45, 46). Single molecule optical force microscopy experiments indicated that upstream sequence could contribute to termination efficiency (47), which was supported by high throughput characterization studies of hundreds of natural termination sites in *E. coli* (35, 48, 49). However, these studies did not demonstrate a mechanism for upstream sequence context to alter termination efficiency, except by potentially misfolding the terminator hairpin itself in a manner analogous to antiterminators. Structural studies of paused RNAP complexes (33, 34) indicated that the RNAP exit channel opening is too narrow to accommodate multiple secondary structures. Relatedly, studies of antitermination factors such as Q, revealed that these factors work by constricting the RNAP exit channel opening to block terminator nucleation (31, 32). We therefore reasoned that nascent RNA itself could be acting as an antiterminator by folding into structures that physically occlude the formation of intrinsic transcriptional terminator hairpins inside the RNAP exit channel.

Using this principle, we designed TOW riboswitches simply by placing aptamer domains next to intrinsic terminator hairpins in such a way that ligand stabilization of the aptamer would physically occlude terminator hairpin formation to cause ligand-mediated antitermination. This design is vastly simpler than all known natural transcriptional riboswitches, which require extensive sequence overlap between aptamer and terminator to create structural competition that determines the gene expression outcome. For example, the folding of the *Cbe pfl* ZTP riboswitch terminator requires competion for 15 nt from the aptamer domain (24). From an engineering perspective, this sequence overlap severely constrains the design space, and requires complex compensatory changes across the molecule if, for example, aptamers are changed (6). While current secondary structure modeling tools have been sufficient to enable the design of translational riboswitches, which primarily operate in a thermodynamic regime (7, 50, 51), the design of transcriptional riboswitches that utilize kinetic switching folding pathways remains limited (52). For this reason, previous attempts to produce synthetic transcriptional riboswitches have attempted to minimize aptamer:expression platform overlap, or have relied on reusing natural switching domains (28, 53, 54). Notably, the TOW mechanism requires no sequence overlap between the aptamer domain and expression platform, enabling modular riboswitch design. We also observe that relatively simple rules can be used to balance the two gene expression outcomes. Termination can be favored by using a stronger (lower ΔG) hairpin, or by extending the single stranded spacer connecting the AD to the EP. Antitermination can be favored by shortening the single-stranded spacer region, or by the addition of ligand-induced stabilizations to the AD. Because of its simpler design, the TOW riboswitch platform is more modular and more easily programmable than previous transcriptional riboswitch systems.

Similar to the discovery of the first natural riboswitches being inspired by success in creating synthetic allosteric ribozymes in the lab (55, 56), the simplicity and success of engineering TOW riboswitches motivated us to search for them in nature. Focusing on the ZTP aptamer, we bioinformatically searched for the TOW riboswitch motif and found 434 candidates out of the 3944 known ZTP riboswitch sequences. Characterizaiton of a subset of these revealed the presence of functional switches, though interestingly the discovered switches mostly reside in *Bacillis*. Further studies comparing the function of switches in *B. subtilis* versus *E. coli* uncovered that nearly all of the switches in the panel are functional in *B. subtilis* and non-functional in *E. coli*, suggesting that there are host-dependent features of TOW antitermination. However, the function of nearly all these riboswitches could be restored in the non-native host by merely strengthening the termination signal, indicating that aptamer folding and ligand-binding functions are retained across large evolutionary distances. At the same time, we also find that NusG plays a nuanced role in facilitating riboswitch folding and/or switching in *B. subtilis*, supporting reports that this transcription factor aids riboswitch function in gram positive bacteria (43, 44). Overall these studies validate that we have discovered the TOW motif to already be present in nature. Due to the simplicity of the motif, we anticipate it to be more broadly distributed and utilized by other riboswitch aptamer families (1). In addition, the simplicity of the motif could motivate its search in higher eukaryotes (57).

In summary, our design of TOW riboswitches represents a system that is simpler, more modular and more easily engineered than previous riboswitch systems, which will enable their expanded use in a broad range of biotechnology applications. Riboswitches are already making impact in a range of important applications in diagnostics (8, 58) and metabolic engineering (59, 60), and the modularity and simplicity of the design rules discovered promise to accelerate the development of robust and tunable systems. Given the availability of many synthetic RNA aptamers that bind small molecules and proteins of diagnostic relevance, we predict the application of high-throughput screening and the development of machine learning tools will enable facile transcriptional riboswitch engineering using the TOW mechanism. Our discovery of TOW riboswitches in nature represents a new mechanism by which nascent RNA structures can regulate transcription and could inform studies of many other nascent RNA structure regulatory mechanisms, for example by RNA binding proteins that bind RNA cotranscriptionally (61), and potentially by the many eukaryotic processes that occur cotranscriptionally and are influenced by nascent RNA structure (62–64).

## MATERIALS & METHODS

### Cloning and plasmid construction

Expression reporter plasmids were prepared as described previously (24). Briefly, reporter plasmids comprised a p15A backbone carrying chloramphenicol resistance, a J23119 *E. coli* σ70 consensus promoter, termination site / riboswitch variant, ribosome binding site, and superfolder green fluorescent protein (sfGFP) coding sequence. Variant sequences were cloned into the backbone either by Gibson assembly using gBlocks (IDT) or inverse PCR using primers (IDT). Plasmid stocks used for data collection were assessed for oligomerization by gel electrophoresis, and sequence-confirmed by Sanger sequencing (Quintara Biosciences). Sequences of all reporter plasmids are available in the Dataset S1.

### B. subtilis strain engineering

Shuttle plasmids were assembled by Gibson assembly using gBlocks (IDT) and the backbone from pBS1C. pBS1C was a gift from Thorsten Mascher (Addgene plasmid #<u>55168</u>) (65). Shuttle plasmids were linearized by BsaI digestion (NEB) and used to transform *B. subtilis* strains PLBS727 (WT) and PLBS728 (*ΔnusG*) grown to stationary phase in MNGE medium following the protocol of Mascher and colleagues (65). PLBS727 (WT) and PLBS728 (*ΔnusG*) were gifts from Paul Babitzke (42). Selected transformants were validated by colony PCR of the amyE locus and assessed for amylase activity by the starch-iodine test. Transformants used for data collection were stored as glycerol stocks at -70ºC. Sequences of all riboswitch reporters are available in Dataset S1.

### In vivo bulk fluorescence reporter assay

*E. coli* reporter expression assays were performed as described previously (24). Briefly, reporter plasmids were transformed into *E. coli* BW25113 (Keio parent strain), and colonies were used to inoculate 300 μL overnight cultures in LB media. Subcultures in M9 enriched media (1x M9 salts, 0.4% glycerol, 0.2% casamino acids, 2 mM MgSO_4_, 0.1 mM CaCl_2_) treated with appropriate antibiotics were inoculated with 4 μL of liquid overnight culture. Subcultures were shaken for 6 hr at 1000 rpm and 37ºC, after which OD_600_ and sfGFP fluorescence were measured using a Biotek Synergy H1 (Agilent) microplate reader operated with BioTek Gen5 (Agilent) software. All in vivo reporter assays were performed with three experimental replicates, each performed in triplicate with three separate colonies (biological replicates) for a total of nine data points (n = 9) per reporter construct.

For riboswitch assays, the indicated amount of ligand was added to the subculture. For ZTP riboswitch assays, 5-aminoimidazole-4-carboxyamide-ribonucleoside (Z) (Millipore Sigma) in DMSO was used; for purine riboswitch assays 2-aminopurine (2AP) (Millipore Sigma) in DMSO was used; for tetracycline riboswitch assays anhydrotetracycline (Cayman Chemical) in ethanol was used. For fluoride riboswitch assays, reporter plasmids were transformed into *E. coli* JW0619 (Δ*crcB* Keio), and subcultures were treated with NaF (Millipore Sigma) in water. Fluorescence data was processed by subtracting the OD_600_ and fluorescence values for blank wells with media only, and normalizing fluorescence values by the OD_600_ value for each sample-containing well.

*B. subtilis* reporter expression assays were performed similarly. Glycerol stocks were used to inoculate overnight cultures in LB, which were used to inoculate subcultures of MNGE media containing the indicated amount of Z. Subcultures were shaken for 6 hr at 37ºC prior to endpoint bulk fluorescence measurements.

### Single round in vitro transcription termination assay

Transcription termination assays were performed as described previously (24). Briefly, 30 second single round IVT reactions were performed with 400 μM NTPs, with reinitiation prevented by the presence of 10 μg/mL rifampicin. Reaction products were isolated by TRIzol-chloroform (Invitrogen) extraction and isolated by TURBO DNase (Invitrogen) digestion followed by a second TRIzol-chloroform (Invitrogen) extraction. Purified IVT products were run on urea polyacrylamide gels, stained with 1x SYBR Gold Nucleic Acid Gel Stain (Invitrogen), and imaged on a Bio-Rad ChemiDoc Touch running Image Lab Touch 3.0. Quantification analysis was performed in Bio-Rad Image Lab 6.1. Bands were added at the expected termination and anti-termination lengths, and manually adjusted to ensure band boundaries were the same across all lanes, and the reported % band intensity of the anti-termination band was recorded. All IVT source gels and quantified data are available in Dataset S1.

### Bioinformatic analysis of ZTP riboswitch expression platforms

ZTP riboswitch expression platform analysis was performed as described previously (66), with some modifications. Briefly, a ZTP riboswitch sequence dataset was prepared by pooling sequences reported by Breaker and colleagues (ref. (18)) with sequences identified in the Rfam database (67) ZTP riboswitch entry <u>RF01750</u> and deduplicating the resulting dataset. Riboswitch sequences were analyzed for the presence of an intrinsic termination site (hairpin with U-rich tract) located downstream of the consensus P3 hairpin sequence. ViennaRNA secondary structure prediction (68) was used to screen for sequences where both the P3 hairpin and terminator hairpin are predicted to form simultaneously (i.e. not mutually exclusive), with a single stranded region in between the two domains. The full list is available in Dataset S1, including the panel selected for functional characterization in vivo. All scripts used for bioinformatic analysis and visualization are available at https://github.com/LucksLab/Bushhouse_Tug-of-War.

## Supporting information

Sequences, Source Data

Source Gel Images

## ACKNOWLEDGEMENTS

We thank Paul Babitzke for sharing the *B. subtilis* strains used in this work, and for helpful discussions about transcription termination in *B. subtilis*. We thank Thorsten Mascher for the shuttle plasmids used to engineer *B. subtilis*, and Irina Artsimovitch for the A26 TSS sequence used to generate several reporter expression plasmids. Support for this work was provided by National Institutes of Health grant 1R35GM161278 to J.B.L., the National Science Foundation Grant 2310382 to J.B.L., the National Institutes of Health Training Grant T32GM008382 through the Northwestern University Molecular Biophysics Training Program to D.Z.B. D.Z.B. was also supported by the Northwestern University Rappaport Award for Research Excellence. J.B.L. was also supported by a John Simon Guggenheim Fellowship. The content is solely the responsibility of the authors and does not necessarily represent the official views of the NIH or NSF.

## AUTHOR CONTRIBUTIONS

Conceptualization, D.Z.B. and J.B.L.; Methodology, D.Z.B. and J.B.L.; Formal analysis, D.Z.B.; Investigation, D.Z.B. and J.F.; Writing–Original Draft, D.Z.B. and J.B.L.; Writing–Review & Editing, D.Z.B., J.F., and J.B.L.; Visualization, D.Z.B.; Software, D.Z.B.; Supervision, J.B.L.; Funding Acquisition, J.B.L.

## DATA AND CODE AVAILABILITY

All source data is provided in Dataset S1. Code is made available at https://github.com/LucksLab/Bushhouse_Tug-of-War.

## REFERENCES

1. P. J. Mccown, K. A. Corbino, S. Stav, M. E. Sherlock, R. R. Breaker, Riboswitch diversity and distribution. RNA 23, 995–1011 (2017).

2. R. R. Breaker, The Biochemical Landscape of Riboswitch Ligands. Biochemistry 61, 137–149 (2022).

3. C. E. Scull, S. S. Dandpat, R. A. Romero, N. G. Walter, Transcriptional Riboswitches Integrate Timescales for Bacterial Gene Expression Control. Front. Mol. Biosci. 7, 607158 (2021).

4. W. Winkler, A. Nahvi, R. R. Breaker, Thiamine derivatives bind messenger RNAs directly to regulate bacterial gene expression. Nature 419, 952–956 (2002).

5. J. A. Howe, et al., Selective small-molecule inhibition of an RNA structural element. Nature 526, 672–677 (2015).

6. J. Hoetzel, T. Wang, B. Suess, Beyond the niche -unlocking the full potential of synthetic riboswitches. Nat Commun 16, 9897 (2025).

7. G. E. Vezeau, L. R. Gadila, H. M. Salis, Automated design of protein-binding riboswitches for sensing human biomarkers in a cell-free expression system. Nat Commun 14, 2416 (2023).

8. W. Thavarajah, et al., Point-of-Use Detection of Environmental Fluoride via a Cell-Free Riboswitch-Based Biosensor. ACSSynBio 9, 10–18 (2019).

9. V. Hedwig, et al., Engineering oxypurinol-responsive riboswitches based on bacterial xanthine aptamers for gene expression control in mammalian cell culture. Nucleic Acids Res 53, gkae1189 (2025).

10. K. Kappel, et al., Accelerated cryo-EM-guided determination of three-dimensional RNA-only structures. Nat Methods 17, 699–707 (2020).

11. K. E. Watters, E. J. Strobel, A. M. Yu, J. T. Lis, J. B. Lucks, Cotranscriptional folding of a riboswitch at nucleotide resolution. NSMB 23, 1124–1131 (2016).

12. K. L. Freida, S. M. Block, Direct Observation of Cotranscriptional Folding in an Adenine Riboswitch. Science 338, 397–400 (2012).

13. A. Chauvier, J. Cabello-Villegas, N. G. Walter, Probing RNA structure and interaction dynamics at the single molecule level. Methods 162–163, 3–11 (2019).

14. A. Reining, et al., Three-state mechanism couples ligand and temperature sensing in riboswitches. Nature 499, 355–359 (2013).

15. L. T. Olenginski, S. F. Spradlin, R. T. Batey, Flipping the script: understanding riboswitches from an alternative perspective. Journal of Biological Chemistry 300, 105730 (2024).

16. M. Mandal, R. R. Breaker, Adenine riboswitches and gene activation by disruption of a transcription terminator. Nat Struct Mol Biol 11, 29–35 (2004).

17. J. L. Baker, et al., Widespread Genetic Switches and Toxicity Resistance Proteins for Fluoride. Science 335, 233–235 (2012).

18. P. B. Kim, J. W. Nelson, R. R. Breaker, An ancient riboswitch class in bacteria regulates purine biosynthesis and one-carbon metabolism. Mol Cell 57, 317–328 (2015).

19. N. White, H. Sadeeshkumar, A. Sun, N. Sudarsan, R. R. Breaker, Lithium-sensing riboswitch classes regulate expression of bacterial cation transporter genes. Sci Rep 12, 19145 (2022).

20. J. K. Wickiser, W. C. Winkler, R. R. Breaker, D. M. Crothers, The speed of RNA transcription and metabolite binding kinetics operate an FMN riboswitch. Mol Cell 18, 49–60 (2005).

21. S. Blouin, R. Chinnappan, D. A. Lafontaine, Folding of the lysine riboswitch: importance of peripheral elements for transcriptional regulation. Nucleic Acids Res 39, 3373–3387 (2011).

22. V. K. Boyapati, W. Huang, J. Spedale, F. Aboul-ela, Basis for ligand discrimination between ON and OFF state riboswitch conformations: The case of the SAM-I riboswitch. RNA 18, 1230–1243 (2012).

23. J. Mulhbacher, D. A. Lafontaine, Ligand recognition determinants of guanine riboswitches. NAR 35, 5568–5580 (2007).

24. D. Z. Bushhouse, J. B. Lucks, Tuning strand displacement kinetics enables programmable ZTP riboswitch dynamic range in vivo. NAR 51, 2891–2903 (2023).

25. L. Cheng, et al., Cotranscriptional RNA strand exchange underlies the gene regulation mechanism in a purine-sensing transcriptional riboswitch. NAR gkac102 (2022). 10.1093/nar/gkac102.

26. E. J. Strobel, L. Cheng, K. E. Berman, P. D. Carlson, J. B. Lucks, A ligand-gated strand displacement mechanism for ZTP riboswitch transcription control. NatureChemBio 15, 1067–1076 (2019).

27. J. E. Weigand, B. Suess, Aptamers and riboswitches: perspectives in biotechnology. Appl Microbiol Biotechnol 85, 229–236 (2009).

28. P. Ceres, J. J. Trausch, R. T. Batey, Engineering modular ‘ON’ RNA switches using biological components. NAR 41, 10449–10461 (2013).

29. S. Dey, et al., Structural insights into RNA-mediated transcription regulation in bacteria. Mol Cell S1097-2765(22)00909–1 (2022). 10.1016/j.molcel.2022.09.020.

30. S. Hwang, et al., Structural basis of transcriptional regulation by a nascent RNA element, HK022 putRNA. Nat Commun 13, 4668 (2022).

31. Z. Yin, J. G. Bird, J. T. Kaelber, B. E. Nickels, R. H. Ebright, In transcription antitermination by Qλ, NusA induces refolding of Qλ to form a nozzle that extends the RNA polymerase RNA-exit channel. PNAS 119, e2205278119 (2022).

32. Z. Yin, J. T. Kaelber, R. H. Ebright, Structural basis of Q-dependent antitermination. Proc Natl Acad Sci U S A 116, 18384–18390 (2019).

33. J. Y. Kang, et al., RNA Polymerase Accommodates a Pause RNA Hairpin by Global Conformational Rearrangements that Prolong Pausing. Molecular Cell 69, 802-815.e5 (2018).

34. L. You, et al., Structural basis for intrinsic transcription termination. Nature 613, 783–789 (2023).

35. Y.-J. Chen, et al., Characterization of 582 natural and synthetic terminators and quantification of their design constraints. NatureMethods 10, 659–664 (2013).

36. L. K. Drogalis, R. T. Batey, Requirements for efficient ligand-gated co-transcriptional switching in designed variants of the B. subtilis pbuE adenine-responsive riboswitch in E. coli. PLoS ONE 15, e0243155 (2020).

37. S. Hanson, K. Berthelot, B. Fink, J. E. G. McCarthy, B. Suess, Tetracycline-aptamer-mediated translational regulation in yeast. Molecular Microbiology 49, 1627–1637 (2003).

38. D. B. Ritchie, D. A. N. Foster, M. T. Woodside, Programmed -1 frameshifting efficiency correlates with RNA pseudoknot conformational plasticity, not resistance to mechanical unfolding. Proceedings of the National Academy of Sciences 109, 16167–16172 (2012).

39. Y. Wang, et al., Comparative studies of frameshifting and nonframeshifting RNA pseudoknots: a mutational and NMR investigation of pseudoknots derived from the bacteriophage T2 gene 32 mRNA and the retroviral gag-pro frameshift site. RNA 8, 981–996 (2002).

40. I. Artsimovitch, V. Svetlov, L. Anthony, R. R. Burgess, RNA Polymerases from Bacillus subtilis and Escherichia coli Differ in Recognition of Regulatory Signals In Vitro. J Bacteriol 182, 6027–6035 (2000).

41. M. J. L. de Hoon, Y. Makita, K. Nakai, S. Miyano, Prediction of Transcriptional Terminators in Bacillus subtilis and Related Species. PLoS Comput Biol 1, e25 (2005).

42. Z. F. Mandell, et al., Comprehensive transcription terminator atlas for Bacillus subtilis. Nat Microbiol 7, 1918–1931 (2022).

43. Z. F. Mandell, D. Zemba, P. Babitzke, Factor-stimulated intrinsic termination: getting by with a little help from some friends. Transcription 13, 96–108 (2022).

44. O. T. Jayasinghe, et al., NusG-dependent RNA polymerase pausing is a common feature of riboswitch regulatory mechanisms. Nucleic Acids Research gkae981 (2024). 10.1093/nar/gkae981.

45. R. Reynolds, R. M. Bermúdez-Cruz, M. J. Chamberlin, Parameters affecting transcription termination by Escherichia coli RNA polymerase. Journal of Molecular Biology 224, 31–51 (1992).

46. R. Reynolds, M. J. Chamberlin, Parameters Affecting Transcription Termination by Escherichia colt’ RNA. Journal of Molecular Biology 224 (1992).

47. M. H. Larson, W. J. Greenleaf, R. Landick, S. M. Block, Applied Force Reveals Mechanistic and Energetic Details of Transcription Termination. Cell 132, 971–982 (2008).

48. G. Cambray, et al., Measurement and modeling of intrinsic transcription terminators. Nucleic Acids Research 41, 5139–5148 (2013).

49. V. K. Mutalik, et al., Precise and reliable gene expression via standard transcription and translation initiation elements. NatureMethods 10, 354–360 (2013).

50. A. Espah Borujeni, D. M. Mishler, J. Wang, W. Huso, H. M. Salis, Automated physics-based design of synthetic riboswitches from diverse RNA aptamers. NAR 44, 1–13 (2016).

51. J. O. L. Andreasson, et al., Crowdsourced RNA design discovers diverse, reversible, efficient, self-contained molecular switches. PNAS 119, e2112979119 (2022).

52. A. Xayaphoummine, V. Vlasnoff, S. Harlepp, H. Isambert, Encoding folding paths of RNA switches. NAR 35, 614–622 (2007).

53. P. Ceres, A. D. Garst, J. G. Marcano-Velázquez, R. T. Batey, Modularity of Select Riboswitch Expression Platforms Enables Facile Engineering of Novel Genetic Regulatory Devices. ACSSynBio 2, 463–472 (2013).

54. S. V. Harbaugh, et al., Engineering a Synthetic Dopamine-Responsive Riboswitch for In Vitro Biosensing. ACSSynBio 11, 2275–2283 (2022).

55. A. Nahvi, et al., Genetic Control by a Metabolite Binding mRNA. Chemistry & Biology 9, 1043–1049 (2002).

56. W. Winkler, A. Nahvi, R. R. Breaker, Thiamine derivatives bind messenger RNAs directly to regulate bacterial gene expression. Nature 419, 952–956 (2002).

57. M. Khoroshkin, et al., A systematic search for RNA structural switches across the human transcriptome. Nat Methods 21, 1634–1645 (2024).

58. W. Thavarajah, et al., The accuracy and usability of point-of-use fluoride biosensors in rural Kenya. NPJ Clean Water 6, 5 (2023).

59. L.-B. Zhou, A.-P. Zeng, Engineering a Lysine-ON Riboswitch for Metabolic Control of Lysine Production in Corynebacterium glutamicum. ACS Synth. Biol. 4, 1335–1340 (2015).

60. L.-B. Zhou, A.-P. Zeng, Exploring Lysine Riboswitch for Metabolic Flux Control and Improvement of l-Lysine Synthesis in Corynebacterium glutamicum. ACS Synth. Biol. 4, 729–734 (2015).

61. M. L. Rodgers, B. O’Brien, S. A. Woodson, Small RNAs and Hfq capture unfolded RNA target sites during transcription. Molecular Cell 83, 1489-1501.e5 (2023).

62. D. Z. Bushhouse, E. K. Choi, L. M. Hertz, J. B. Lucks, How does RNA fold dynamically? JMB 434, 167665 (2022).

63. Y. Yoon, C. Quan, L. V. Soles, Y. Shi, Coordinating mRNA maturation: The U1 relay model. Molecular Cell 86, 449–460 (2026).

64. M. Shine, et al., Co-transcriptional gene regulation in eukaryotes and prokaryotes. Nat Rev Mol Cell Biol 25, 534–554 (2024).

65. J. Radeck, et al., The Bacillus BioBrick Box: generation and evaluation of essential genetic building blocks for standardized work with Bacillus subtilis. Journal of Biological Engineering 7, 29 (2013).

66. D. Z. Bushhouse, J. Fu, J. B. Lucks, RNA folding kinetics control riboswitch sensitivity in vivo. Nat Commun 16, 953 (2025).

67. N. Ontiveros-Palacios, et al., Rfam 15: RNA families database in 2025. Nucleic Acids Research 53, D258–D267 (2025).

68. R. Lorenz, et al., ViennaRNA Package 2.0. Algorithms Mol Biol 6, 26 (2011).

69. A. Ren, K. R. Rajashankar, D. J. Patel, Fluoride ion encapsulation by Mg2+ ions and phosphates in a fluoride riboswitch. Nature 486, 85–89 (2012).

70. J. R. Stagno, et al., Structures of riboswitch RNA reaction states by mix-and-inject XFEL serial crystallography. Nature 541, 242–246 (2017).

71. H. Xiao, T. E. Edwards, A. R. Ferré-D’Amaré, Structural Basis for Specific, High-Affinity Tetracycline Binding by an In Vitro Evolved Aptamer and Artificial Riboswitch. Chemistry & Biology 15, 1125–1137 (2008).

