## Supplementary Figures for "Design and discovery of ‘tug-of-war’ riboswitches"

### Supporting Information for
Design and discovery of ‘tug-of-war’ riboswitches

David Z. Bushhouse, Jiayu Fu, & Julius B. Lucks

Julius B. Lucks


**This PDF file includes:**

Figures S1 to S4
Legend for Dataset S1

**Other supporting materials for this manuscript include the following:**

Dataset S1

#### Figures

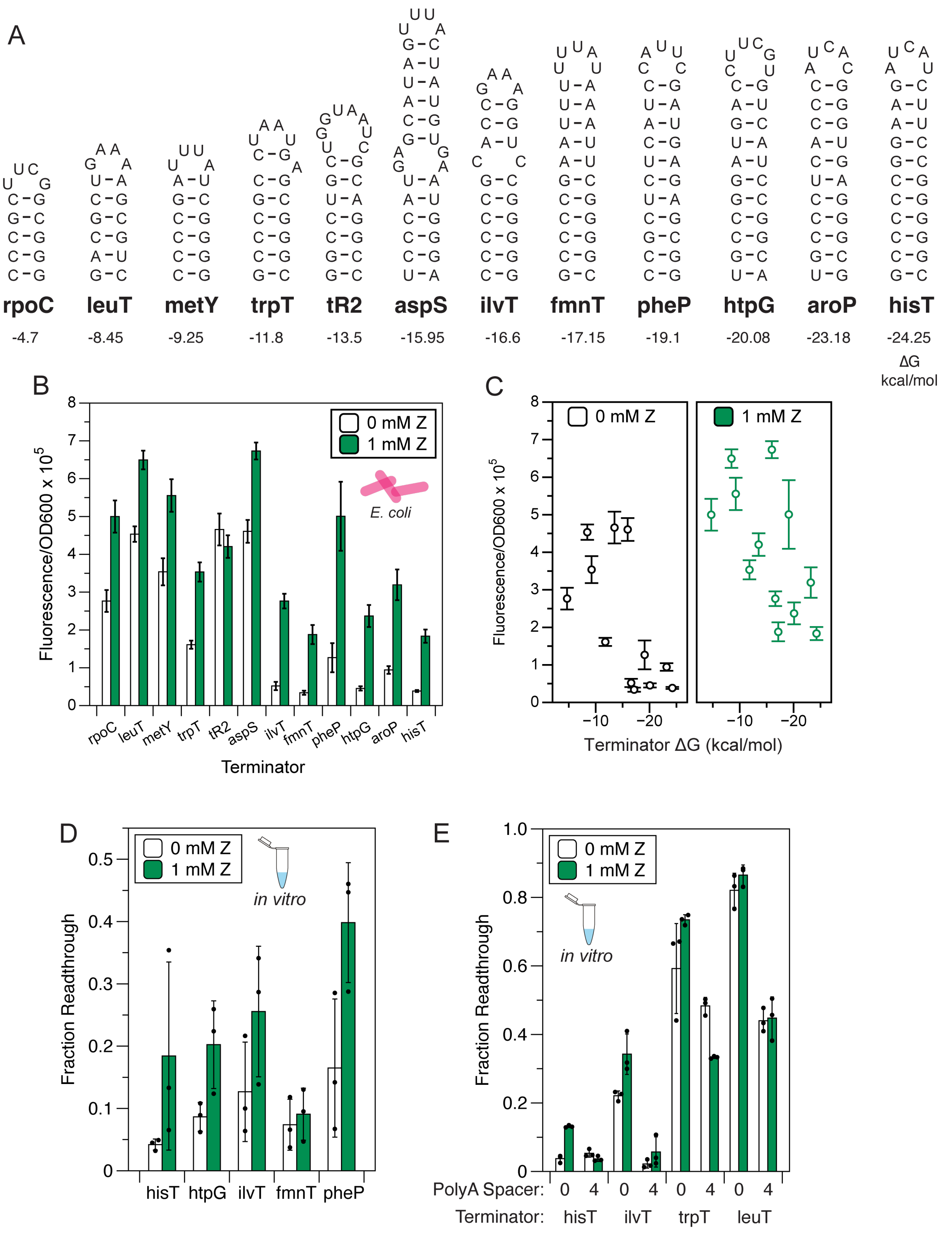


**Fig. S1. Cellular and in vitro characterization of synthetic TOW ZTP riboswitches.** (A) NUPACK secondary structure predictions of the terminator panel used to construct synthetic TOW ZTP riboswitches according to the design of Figure 1B. (B) In vivo reporter expression assays performed in E. coli characterizing switching function of TOW riboswitch designs with 0 mM Z and

1 mM Z treatment. Error bars indicate standard deviation of n=9 biological replicates. (C) Reporter expression results from (B) plotted against the NUPACK predicted $\Delta G$ of the terminator stem. Error bars indicate standard deviation of n=9 biological replicates. (D) IVT termination assays performed on a subset of the constructs assessed in (B). (E) IVT antitermination assays performed on a subset of the constructs assessed in Fig. 1C. For (D-E), points indicate quantified fraction antitermination, bars indicate mean, and error bars indicate standard deviation over n=3 replicates. Source gels are available in Dataset S1.

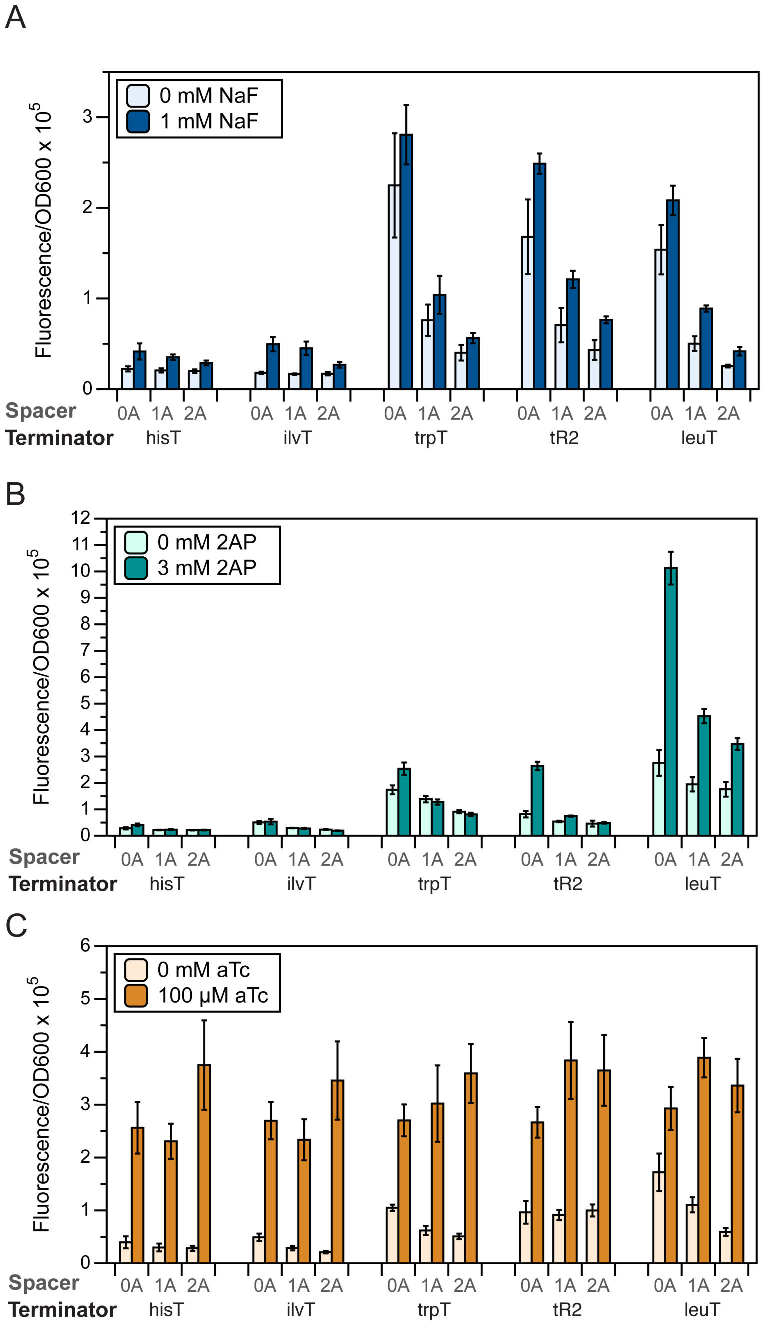


**Fig. S2. Functional characterization of synthetic TOW riboswitches.** (A) In vivo reporter expression characterization of synthetic TOW fluoride riboswitch designs in the presence (1 mM NaF) and absence (0 mM NaF) of cognate ligand. (B) In vivo reporter expression characterization of synthetic TOW purine riboswitch designs in the presence (3 mM 2-AP) and absence (0 mM 2-AP) of cognate ligand. (C) In vivo reporter expression characterization of synthetic TOW tetracycline riboswitch designs in the presence (100 uM aTc) and absence (0 uM aTc) of cognate ligand. For A-C error bars indicate standard deviation of n=9 biological replicates. Designs were constructed according to the schematics in Figure 2 B, E, H.

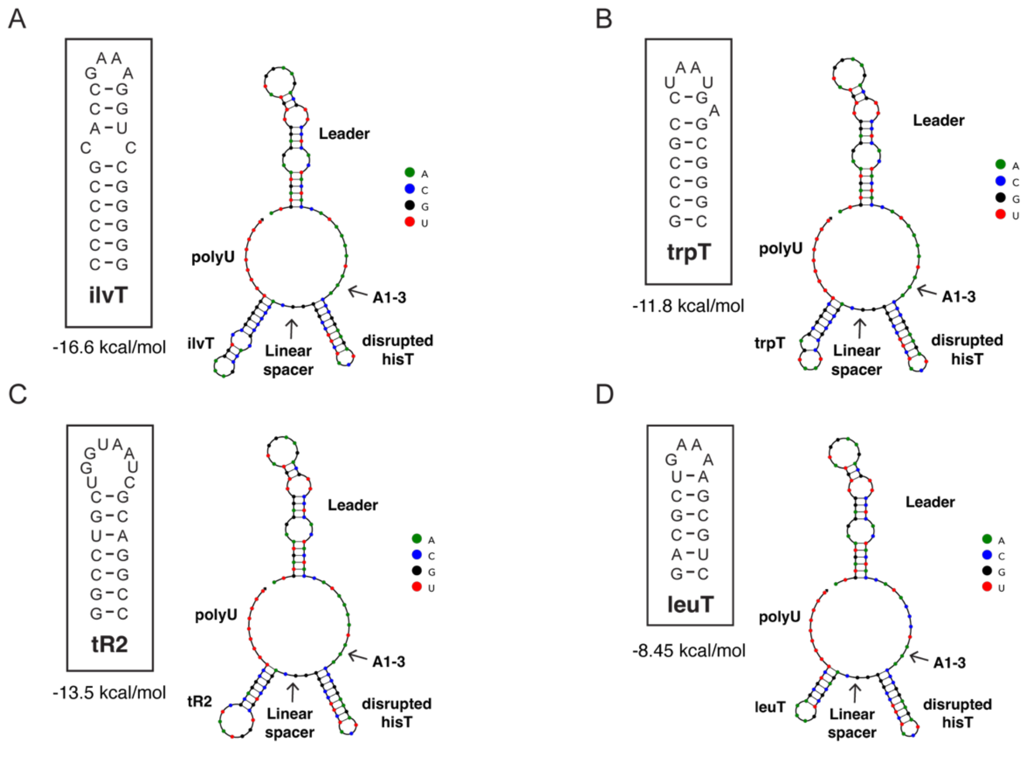


**Fig. S3. Design of reporter expression assay hisT stem disruption variants from Fig. 3C.** (A) NUPACK secondary structure prediction of the hisT(3A)-A-ilvT variant in which the 3 5' bases of the hisT stem have been mutated to A. The disruption of the hisT stem is predicted to form a linear region between the shortened hisT stem and the intact ilvT stem. Inset depicts the NUPACK secondary structure prediction of the ilvT stem alone, and the predicted ΔG at 37 °C. (B) As in (A) for the hisT(3A)-A-trpT variant in which the 3 5' bases of the hisT stem have been mutated to A. (C) As in (A) for the hisT(3A)-A-tR2 variant in which the 3 5' bases of the hisT stem have been mutated to A. (D) As in (A) for the hisT(3A)-A-leuT variant in which the 3 5' bases of the hisT stem have been mutated to A.

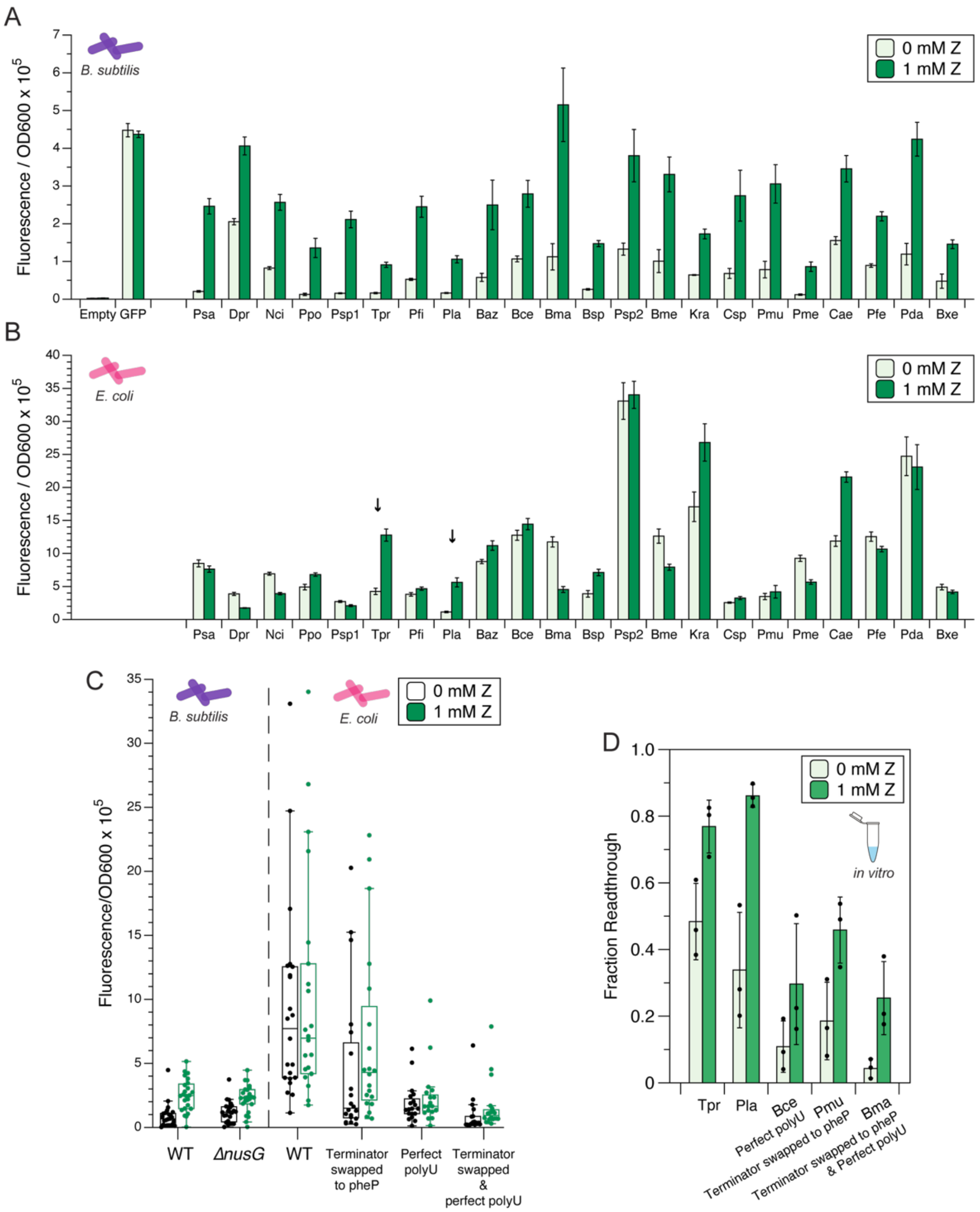


**Figure S4. Discovery of natural ZTP riboswitches with functional TOW architecture.** (A) In vivo reporter expression characterization of natural ZTP riboswitch panel in the presence (1 mM Z) and absence (0 mM Z) of ligand in *B. subtilis*. Error bars indicate standard deviation of n=9 biological replicates. (B) In vivo reporter expression characterization of natural ZTP riboswitch panel in the presence (1 mM Z) and absence (0 mM Z) of ligand in *E. coli*. Error bars indicate standard deviation of n=9 biological replicates. (C) Box plots of the terminator readthrough expression fluorescence with 0 mM Z and 1 mM Z treatment for riboswitches from the experimental panel. Points indicate mean fluorescence values for each riboswitch. Data left of the dotted line was collected from experiments performed in *B. subtilis* stain PLBS727 and PLBS728 as described in Methods. Data right of the dotted line was collected from experiments performed in *E. coli* BW25113 Keio parent strain. X-axis labels on the right side of the plot indicate WT or modified sequences. (D) In vitro transcription assay characterization of Tpr and Pla riboswitches, and select riboswitches with rescued switching function (>2x fold change) in *E. coli*. n = 4, points indicate quantified fraction antitermination, bars indicate mean, and error bars indicate standard deviation. Source gels are available in Dataset S1.

**Dataset S1 (separate file).** Contains sequences of non-riboswitch sequence components, list of all plasmids and strains used in this study, with the sequence of their respective riboswitch inserts, all in vivo fluorescence data, all IVT data, all source gels, and natural TOW ZTP riboswitches identified bioinformatically, including panel selected for characterization.
